# Do monkeys see the way we do? Qualitative similarities and differences between monkey and human perception

**DOI:** 10.1101/2025.01.30.635614

**Authors:** Thomas Cherian, Georgin Jacob, S.P. Arun

**Affiliations:** Centre for Neuroscience, Bengaluru, India; Department of Electrical Communication Engineering, Indian Institute of Science, Bengaluru, India

## Abstract

Monkeys are widely used as models for human vision. While their anatomy and physiology are similar to humans, it is unclear to what extent their perception matches with ours. Previous studies on monkeys have evaluated specific perceptual phenomena after extensive training that could have even altered perception. To resolve these issues, we trained monkeys to perform a highly general oddball visual search task on natural images, and then tested them on carefully controlled image sets to detect a variety of perceptual phenomena and compared their performance with humans. This revealed a number of qualitative similarities and differences. Like humans, monkeys showed similar object relations, Weber’s law and amodal completion. However, unlike humans, monkeys did not show mirror confusion or a global advantage. Our results represent a first comprehensive comparison of visual perception in monkeys and humans, revealing both strengths and limitations of monkeys as a model organism for human visual perception.

## INTRODUCTION

> *The Master sees things as they are, without trying to control them. -- Tao Te Ching* (Mitchell, 1988)

Macaque monkeys are widely used as model organisms to study the neural basis of cognition (Passingham, 2009; Roelfsema and Treue, 2014; Buffalo et al., 2019). Specifically in the case of vision, monkeys have similar visual acuity and contrast thresholds (Weinstein and Grether, 1940; De Valois et al., 1974b, 1974a), and have extensively characterized visual regions whose homologues continue to be elaborated and studied in humans (Quiroga et al., 2005; Hiramatsu et al., 2011; Tang et al., 2014). Moreover, humans and monkeys share similar object representations at a global level (Kriegeskorte et al., 2008a; Rajalingham et al., 2015, 2018), and their visual cortexes also show similar segregation into functional sub-units (Bell et al., 2009). These converging lines of evidence confirm that macaque monkeys are indeed an excellent model for human visual cognition.

However, our understanding of any model is incomplete without knowing its limitations. In this case, it is not clear if there are any fundamental differences in visual perception between monkeys and humans. Do monkeys see the way we do? A comprehensive answer to this question would require comparing monkeys and humans directly on a battery of tests of visual perception. However, this can be extremely challenging in practice. Humans can easily be instructed verbally to perform a variety of visual tasks, but monkeys require extensive behavioural training on each specific visual task. As a result, most studies have tested primates on various specific aspects of perception, as detailed below.

At a coarse level, monkeys trained on categorization tasks show similar performance on natural objects as humans (Rajalingham et al., 2015, 2018). They also show similar gaze patterns on normal faces and illusory faces (Dahl et al., 2010; Weldon et al., 2013; Beran et al., 2017; Taubert et al., 2017). But on other measures, humans and monkeys have shown different patterns of performance. While humans are faster to report shape at the global level compared to the local level (Navon, 1977; Kimchi, 1992), there is conflicting evidence for this effect in monkeys: monkeys trained on shape categorization responded faster to global shape (Tanaka and Fujita, 2000), but in other cases responded faster to local shape (Deruelle and Fagot, 1997; Fagot and Tomonaga, 1999; Nielsen et al., 2006). These differences could arise due to task or stimulus differences.

At a more fine-grained level, macaque monkeys have been tested on a variety of visual illusions. Monkeys trained on size matching tasks are susceptible to size illusions like we are (Behar and Samuel, 1982; Fujita, 1996, 1997; Barbet and Fagot, 2002; Parron and Fagot, 2007; Agrillo et al., 2019), although they are less sensitive to context than humans (Fujita, 1997). Monkeys trained on length-matching tasks show the Muller-Lyer illusion (Tudusciuc and Nieder, 2010). Monkeys trained on image matching tasks show mirror confusion on abstract shapes (Riopelle et al., 1964; Washburn, 1993). Monkeys trained on numerosity tasks show a Weber’s law for numerosity (Nieder and Miller, 2003; Jordan and Brannon, 2006) and biased responses to regular dot configurations (Beran, 2006). However, they do not show certain grouping illusions (Parrish et al., 2019).

### Overview of this study

A major limitation of the above studies is that each finding is based on a highly specialized training regimen tested on a different group of individuals. As a result, we lack a comprehensive evaluation of visual perception in monkeys and how this might compare to humans. Here, we overcame this limitation by training monkeys on a generic oddball visual search task, evaluating multiple perceptual phenomena within the same overall task and comparing these patterns of performance to humans.

In the oddball search task, the participant has to touch the oddball item in an array containing one oddball item and multiple identical distractors (Figure 1). This is an extremely easy task for human participants and required minimal training for monkeys to reach high levels of performance (see Methods). Furthermore, the inverse of the response time of an oddball search is a natural index of the dissimilarity between the target and distractors (Arun, 2012; Pramod and Arun, 2014, 2016a; Sunder and Arun, 2016). The reciprocal of the reaction time also renders the distribution of response times to be less skewed and more normal (Carpenter et al., 2009; Jacob and Arun, 2020). Together, this makes the reciprocal of response time in visual search an objective measure of visual dissimilarity to compare between monkeys and humans.

**Figure 1.**
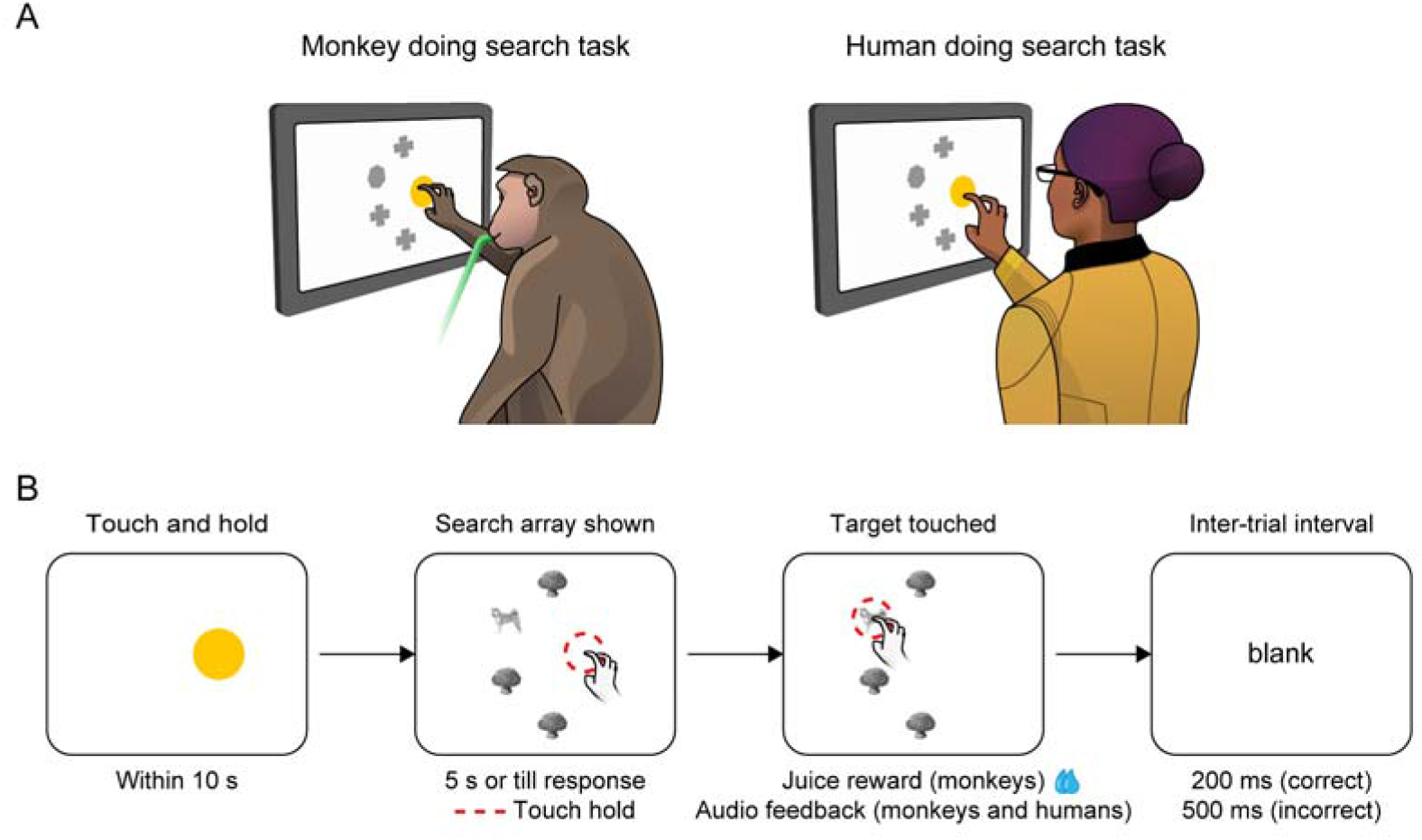
Testing visual perception in monkeys and humans. (A) Schematic of the setup used to test visual perception in monkeys and humans. Monkeys were given a juice reward on making a correct response (indicated by the green juice pipe). An example video of one of the monkeys performing this task is available in <u>Movie S1</u>. Humans received no such tangible rewards but were instead compensated monetarily for their participation. (B) Schematic of the oddball visual search task used to test visual perception in monkeys and humans. Participants had to touch and hold (red dashed lines are for our reference) a yellow button till a search array appeared. The search array always contained one oddball item among multiple identical distractors. Participants could touch any item to complete the trial; touching the oddball target was taken as a correct response.

Importantly, oddball search can be used to measure a variety of perceptual phenomena depending on the specific images used (Jacob et al., 2021b). For example, it is well known that humans experience confusion between vertical mirror images. In visual search, mirror confusion is evidenced by longer response times for finding a target among distractors when the target is a vertical mirror image of the distractor than when it is a horizontal mirror image. Likewise, according to the global advantage effect, humans can report global shape faster than local shape. In visual search, the global advantage effect manifests as faster response times for targets differing in global shape compared to when they differ in local shape.

We identified several perceptual phenomena for testing on both humans and monkeys, which included those used recently to compare humans with deep neural networks (Jacob et al., 2021b). These phenomena all involve comparing closely related pairs of images (e.g. horizontal vs vertical mirror images) where the perceptual effect in most cases cannot be explained using pixel or low-level differences alone. These phenomena included testing coarse object similarities, Weber’s law for luminance, amodal completion of occluded objects, mirror-image confusion, and global advantage. These revealed both qualitative similarities as well as differences between human and monkey perception.

### Movie S1. Video of a monkey performing a visual search task

The video shows monkey M1 performing a visual search task involving images from Experiments 4 & 5 (global & local processing)

## RESULTS

To obtain a comprehensive comparison of visual perception in monkeys and humans, we tested 3 monkeys and 24 humans on an oddball visual search task (Figure 1) across several experiments. Each experiment involved a different set of images designed to address a specific aspect of visual processing, but importantly, the task remained unchanged for the participants: they had to always locate an oddball item in the search array. This design ensured that task considerations of the participants (humans and monkeys) were completely unrelated to the goals of each experiment.

### Experiment 1: Do monkeys have object representations similar to humans?

In Experiment 1, we asked whether monkeys and humans perceive objects similarly at the coarse level. Since visual search is more difficult when the target is similar to the distractors, the reciprocal of the response time can be taken as a measure of perceived dissimilarity between the target and distractor. We therefore asked how well the perceived dissimilarities between objects for monkeys matched those obtained from humans.

To this end, we chose a set of animate and inanimate objects (Figure 2A) and created search arrays involving all possible pairs of objects. We analysed previously published data from 12 human participants on an oddball search task (see Methods). Human reaction times were highly consistent (*r_sh_* = 0.90, *p_sh_* < 0.005, *n* = 28 search pairs; see Methods). To calculate perceived dissimilarity, we took the reciprocal of the average response time (across trials and participants) for each image-pair. The resulting pairwise dissimilarities are depicted in Figure 2B. It can be seen that humans perceive animate objects to be similar to each other, and dissimilar to inanimate objects, as evidenced by the blue cluster in Figure 2B.

**Figure 2.**
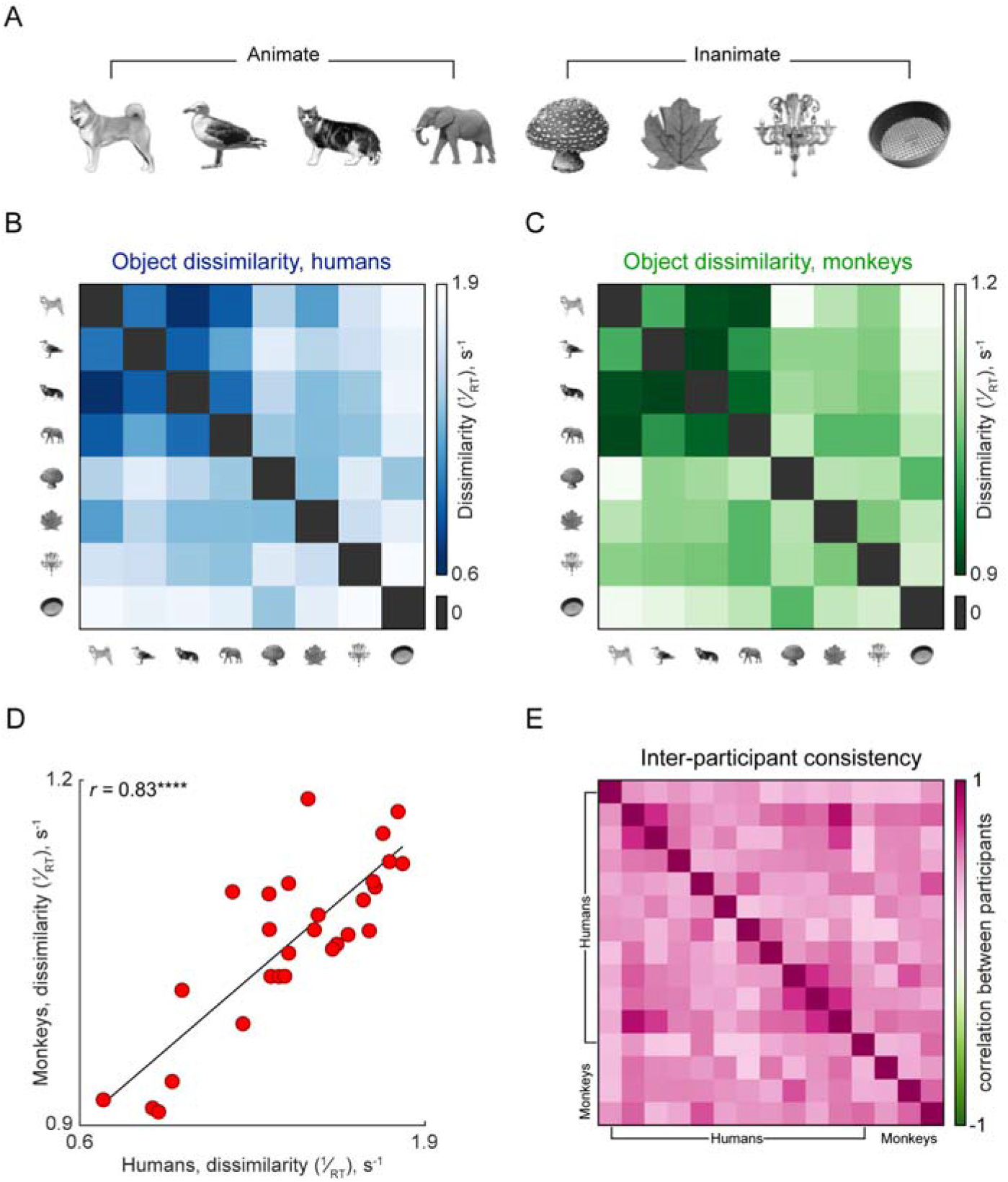
Monkeys have similar coarse object representations as humans. (A) Animate and inanimate objects used in Experiment 1. (B) Object dissimilarities measured from visual search for humans, who perceive animate objects to be similar to each other (blue cluster), and dissimilar to other objects. (C) Same as in (B), but for monkeys who also perceive animate objects to be similar to each other (green cluster) and dissimilar to all other objects. (D) Object dissimilarities for monkeys plotted against object dissimilarities in humans across all 28 pairs of objects tested. *r* is the Pearson’s correlation (**** indicates *p* < 0.00005). (E) Inter-participant consistency for all pairs of participants (both human and monkey), calculated as the pairwise correlation between participants’ object dissimilarities. Large values indicate high agreement between that pair of subjects.

We then proceeded to ask if monkeys show similar object representations as humans. All three monkeys were highly accurate in this task (mean ± s.d.: 82% ± 5%; note that chance accuracy is 25% since there are four items in each search array) and their reaction times were consistent (*r_sh_* = 0.82, *p_sh_*< 0.00005, *n* = 28 search pairs). We then calculated pairwise dissimilarities as before, and the resulting dissimilarity matrix is depicted in Figure 2C. Here too, we find that monkeys perceived animate objects to be similar to each other and dissimilar to inanimate objects (Figure 2C, green cluster).

To quantify the overall match between perceived object dissimilarities between monkeys and humans, we plotted the dissimilarity of each pair of objects for monkeys against the corresponding value obtained from humans (Figure 2D). This revealed a strong and statistically significant correlation (Figure 2D; *r* = 0.83, *p* < 0.00005, *n* = 28 search pairs).

The global match in perceived dissimilarities between humans and monkeys could potentially obscure individual differences. To evaluate this possibility, we calculated the pairwise object dissimilarity for each participant (monkey or human) and calculated the correlation between these dissimilarities for every pair of participants. A high correlation implies that searches that were easy for one participant were also easy for the other, which implies high agreement. Conversely, a low correlation could arise if searches that are easy for one participant are hard for the other. If monkeys and humans had qualitatively different object perception, we would expect them to form two groups, where monkeys’ responses agree with each other but not with humans and likewise human responses agree with each other more than with monkeys. However, we observed no such pattern (Figure 2E), suggesting that there is a good overall match in object perception between monkeys and humans.

We conclude that monkeys have similar object representations as humans at least at a coarse level.

### Experiment 2: Do monkeys show Weber’s law like humans?

Next, we tested humans and monkeys on a universal law of psychophysics, namely Weber’s law. According to Weber’s law, the just noticeable difference in a stimulus depends on the baseline magnitude of the stimulus. Weber’s law is interesting because it is not a low-level visual property but rather requires higher order processing of relative rather than absolute magnitude. We selected luminance as a simple visual feature and asked whether visual search for oddball targets differing in luminance would follow Weber’s law.

To this end, we created search displays in which the oddball target differed only in luminance from the distractors (Figure 3A). According to Weber’s law, participants should find it easy to detect the same luminance change when the distractors have low luminance than when they have high luminance. Conversely, the difficulty of search should be similar when the percentage luminance change is the same, regardless of the absolute luminance. We tested these key predictions in human and monkey participants.

**Figure 3.**
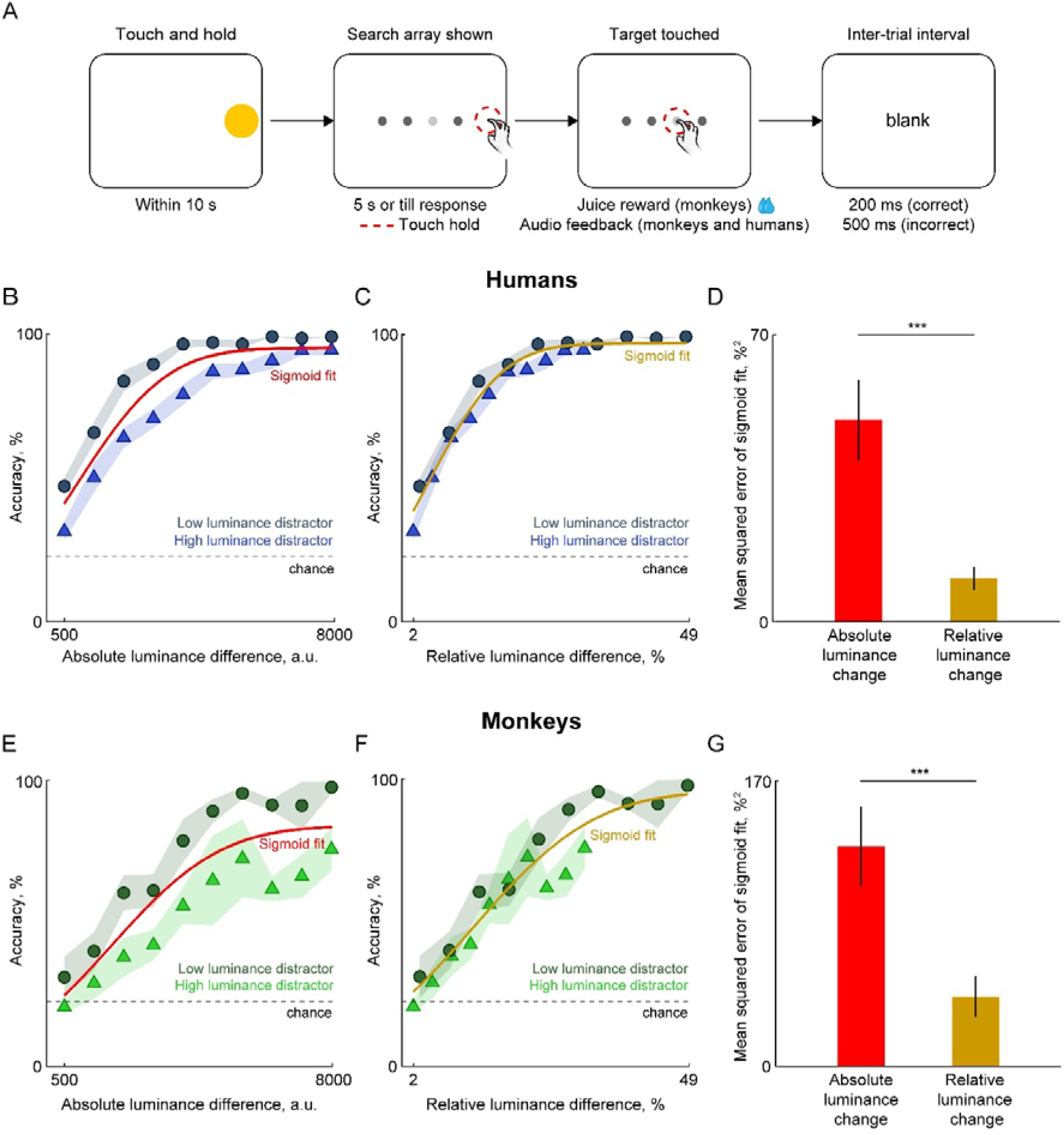
Monkeys experience Weber’s law like humans. (A) Schematic of an oddball visual search trial for testing luminance discrimination. The stimuli were placed in a line centred on the screen and participants had to touch the brightest stimuli, which was always the target, to complete the trial successfully. (B) Accuracy of humans on oddball search for targets differing in luminance as a function of absolute luminance change for high luminance (*blue*) or low luminance distractors (*gray*). Shaded regions depict s.e.m. of accuracy across 16 participants. The best-fitting sigmoid line across both conditions is overlaid in *red*. The dotted line indicates chance accuracy level. (C) Same data as in panel B but replotted as a function of percent change in luminance of the target relative to the distractor. The best fitting sigmoid line across both conditions is overlaid in *ochre*. The dotted line indicates chance accuracy level. (D) Mean squared error between the sigmoid fit and the observed data for absolute luminance changes (*red*) and relative luminance changes (*ochre*). Asterisks indicate statistical significance obtained from a rank-sum test across 20 accuracy values in both conditions (*** indicates *p* < 0.0005). (E-G) Same as B-D but for data from 3 monkeys doing the same task.

Humans were highly accurate on this task (mean ± s.d.: 82% ± 5%) and their reaction times were also highly consistent (*r_sh_* = 0.97, *p_sh_* < 0.005, *n* = 20 search pairs). The performance of 16 human participants is shown in Figure 3B-D. When we plotted accuracy as a function of absolute luminance changes, humans showed greater accuracy for detecting larger luminance changes (Figure 3B). But this accuracy was larger for low-luminance distractors (absolute luminance = 16,384 a.u.) compared to high-luminance distractors (absolute luminance = 25,600 a.u.). Importantly, these accuracy differences disappeared when the same data was replotted as a function of percentage difference in luminance (Figure 3C), which is consistent with Weber’s law.

To quantify these differences, we fit a sigmoid function to the mean accuracy values for both the low-luminance and high-luminance distractor conditions, and compared the mean squared error between the observed accuracies and the sigmoid fits when the data was plotted as a function of absolute luminance difference and when it was plotted as a function of relative changes in luminance. We found a smaller mean-squared error for relative intensity differences (Figure 3D; squared error in accuracy, mean ± s.e.m.: 49.18 ± 9.85 %^2^ for absolute luminance differences; 10.56 ± 2.85 %^2^ for relative luminance differences, *p* < 0.0005, rank-sum test, *n* = 20 search pairs). This confirms that humans showed Weber’s law for detecting luminance differences.

Next, we asked whether monkeys also show similar sensitivity to relative luminance compared to absolute luminance. Monkeys also performed well above chance overall (accuracy, mean ± s.d.: 64% ± 4%) and their reaction times were also consistent (*r_sh_* = 0.69, *p_sh_* < 0.05, *n* = 20 search pairs). However, they showed a shallower increase in accuracy with absolute luminance changes compared to humans (Figure 3E). Interestingly, they too showed greater accuracy for detecting luminance changes in low-luminance distractors compared to high-luminance distractors (Figure 3F). This difference in accuracy reduced greatly when the same data was replotted using relative changes in luminance (Figure 3G). This was evident upon comparing the mean-squared error in the sigmoid fits for absolute and relative luminance differences (Figure 3F; squared error, mean ± s.e.m.: 131.09 ± 23.78 %^2^ for absolute luminance differences; 41.46 ± 12.09 %^2^ for relative luminance differences, *p* < 0.0005, rank-sum test, *n* = 20 search pairs).

To confirm that these differences are present in individual participants, we repeated these analyses for each participant separately. Nearly all (15/16) human participants and all 3 monkey participants showed a larger error in the sigmoid fit for absolute luminance changes compared to relative luminance changes.

We conclude that monkeys, like humans, experience Weber’s law for detecting luminance changes in visual search.

### Experiment 3. Do monkeys perceive occluded shapes like humans?

When we see an occluded display such as the one on the left of Figure 4A, we have no trouble understanding the shape that continues behind the occluder. This kind of amodal completion is thought to be rapid and automatic based on visual search studies (Rensink and Enns, 1998). In these studies, humans find it hard to locate an occluded display among distractors with “likely completion” compared to finding a “mosaic” display in which the two shapes abut without occluding (Figure 4A). Thus, the dissimilarity between occluded and likely displays is smaller than the dissimilarity between likely and mosaic displays. We set out to investigate if monkeys also experience amodal completion as humans do.

**Figure 4.**
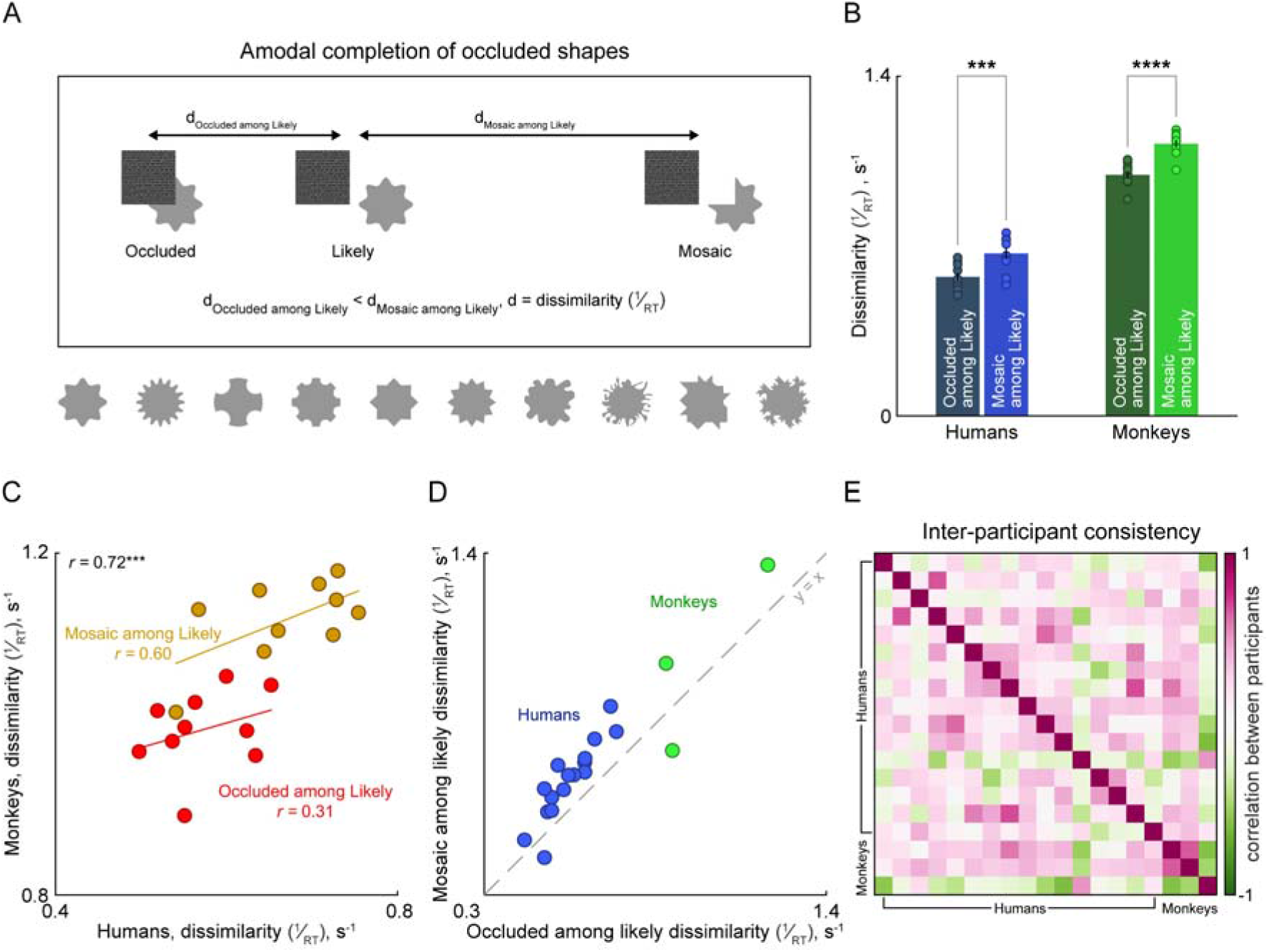
Monkeys experience rapid amodal completion like humans. (A) Occluded objects are rapidly and automatically completed during visual search. In the key comparison, search for an occluded display (*left*) among “likely completions” (*middle*) as distractors is hard, implying that the two displays have low dissimilarity. By contrast, searching for a “mosaic” display (right) among likely completions (*middle*) is easy, implying that these two displays are dissimilar. The shapes at the bottom show the full set of stimuli tested in the experiment. (B) Average dissimilarities for occluded-likely and mosaic-likely searches for humans (*blue, left*) and monkeys (*green, right*). Asterisks indicate statistical significance using a t-test across the dissimilarities of 10 shapes tested (*** is *p* < 0.0005; **** is *p* < 0.00005). (C) *Effect strength for each condition.* Dissimilarities in visual search for monkeys across all searches tested plotted against the corresponding values for humans. The correlation between all points is indicated on the top left (*r* = 0.72). Correlations for each group of pairs are also shown separately: occluded among likely (*red dots*) and mosaic among likely (*ochre dots*). Asterisks represent statistical significance (*** is *p* < 0.0005). Correlation values without an asterisk are not statistically significant. (D) *Effect strength for each participant.* Average dissimilarities of each participant for occluded-likely searches plotted against the dissimilarities for the corresponding mosaic-likely searches. Blue dots represent the 16 human participants; green dots represent 3 monkeys. The dashed line indicates the y = x line. It can be seen that nearly all human and monkey participants showed a larger dissimilarity for mosaic-likely searches compared to occluded-likely searches, confirming the amodal completion effect. (E) Inter-participant consistency for all pairs of participants (both human and monkey), calculated as the pairwise correlation between participants’ object dissimilarities. Large values indicate high agreement between that pair of subjects.

To test amodal completion in monkeys, we took 10 shapes (Figure 4A) and their occluded, likely and mosaic versions from a previous study (Cherian and Arun, 2024). For each shape we used the likely display as the distractor with the occluded or mosaic display as the targets. We then compared the dissimilarities measured in monkeys to humans on the same search pairs.

Humans were highly accurate on this task (accuracy, mean ± s.d.: 96 ± 4% across 16 participants; chance is 8.3%) and their reaction times were consistent (*r_sh_* = 0.64, *p_sh_*= 0.07, *n* = 20 search pairs). We calculated perceived dissimilarity in visual search as before, by taking the reciprocal of the average response time in each condition. In humans, the dissimilarity between occluded and likely displays were significantly less than the dissimilarity between mosaic and likely displays (Figure 4B; dissimilarity, mean ± s.e.m.: 0.57 ± 0.16 s^-1^ for occluded targets among likely displays, 0.66 ± 0.23 s^-1^ for mosaic targets among likely displays; *p* < 0.0005, paired t-test across 10 searches). This confirms that humans experience amodal completion.

Monkeys also performed this task accurately (accuracy, mean ± s.d.: 75% ± 12% across three monkeys; chance is 25%) and their reaction times were consistent (*r_sh_* = 0.72, *p_sh_*= 0.001, *n* = 20 search pairs). Even for monkeys, the dissimilarity between occluded and likely displays was significantly smaller than the dissimilarity between likely and mosaic displays (Figure 4B; dissimilarities, mean ± s.e.m.; 0.99 ± 0.14 s^-1^ for occluded among likely displays; 1.12 ± 0.15 s^-1^ for mosaic among likely displays; *p* < 0.00005, paired t-test across 10 searches of each type).

To quantify the overall match between perceived dissimilarities between monkeys and humans, we plotted the dissimilarity for each search condition for monkeys against the corresponding value for humans. This revealed a strong and statistically significant correlation (Figure 4C; *r* = 0.72, *p* < 0.0005). Mosaic among Likely dissimilarities were correlated between monkeys and humans although this did not reach statistical significance (Figure 4C; *r* = 0.60, *p* = 0.06). Finally, Occluded among Likely dissimilarities were not correlated (Figure 4C; *r* = 0.31, *p* = 0.37).

We then asked if these trends in dissimilarity hold at the individual level. To this end, we plotted the average dissimilarity for mosaic among likely searches for each participant against the average dissimilarity for occluded among likely searches (Figure 4D). Values above the y = x line would confirm the amodal completion effect. Nearly all individual participants, except one human and one monkey, showed the effect (Figure 4D). To further compare individual monkey and human participants, we computed a similarity score between pairs of participants as the correlation between the dissimilarities across the searches. The resulting pairwise dissimilarities are shown in Figure 4E. This revealed no clear grouping of monkey or human participants, again underscoring the fact that monkeys show very similar responses to amodal completion as humans.

We conclude that monkeys experience amodal completion in visual search, just like humans.

### Experiment 4. Do monkeys show mirror confusion like humans do?

Humans tend to confuse vertical mirror images compared to horizontal mirror images, presumably because we frequently see vertical images across changes in viewpoint (Rollenhagen and Olson, 2000; Corballis, 2018). Mirror confusion has been observed in high-level visual cortex in monkeys (Rollenhagen and Olson, 2000) as well as in humans (Corballis, 2018). However, there has been only two studies showing that macaque monkeys show low accuracy in learning to discriminate vertical mirror images of artificial shapes (Riopelle et al., 1964; Washburn, 1993).

We sought to systematically investigate this issue by asking if humans and monkeys experience mirror confusion in visual search involving natural images. We selected 15 natural objects and created horizontal and vertical mirror images of each (Figure 5A). If participants experience vertical mirror confusion, they should be slower to respond on trials when the target is a vertical mirror image compared to when the target is a horizontal image. In terms of dissimilarity, the dissimilarity between vertical mirror images should be smaller than the dissimilarity between horizontal mirror images (Figure 5A).

**Figure 5.**
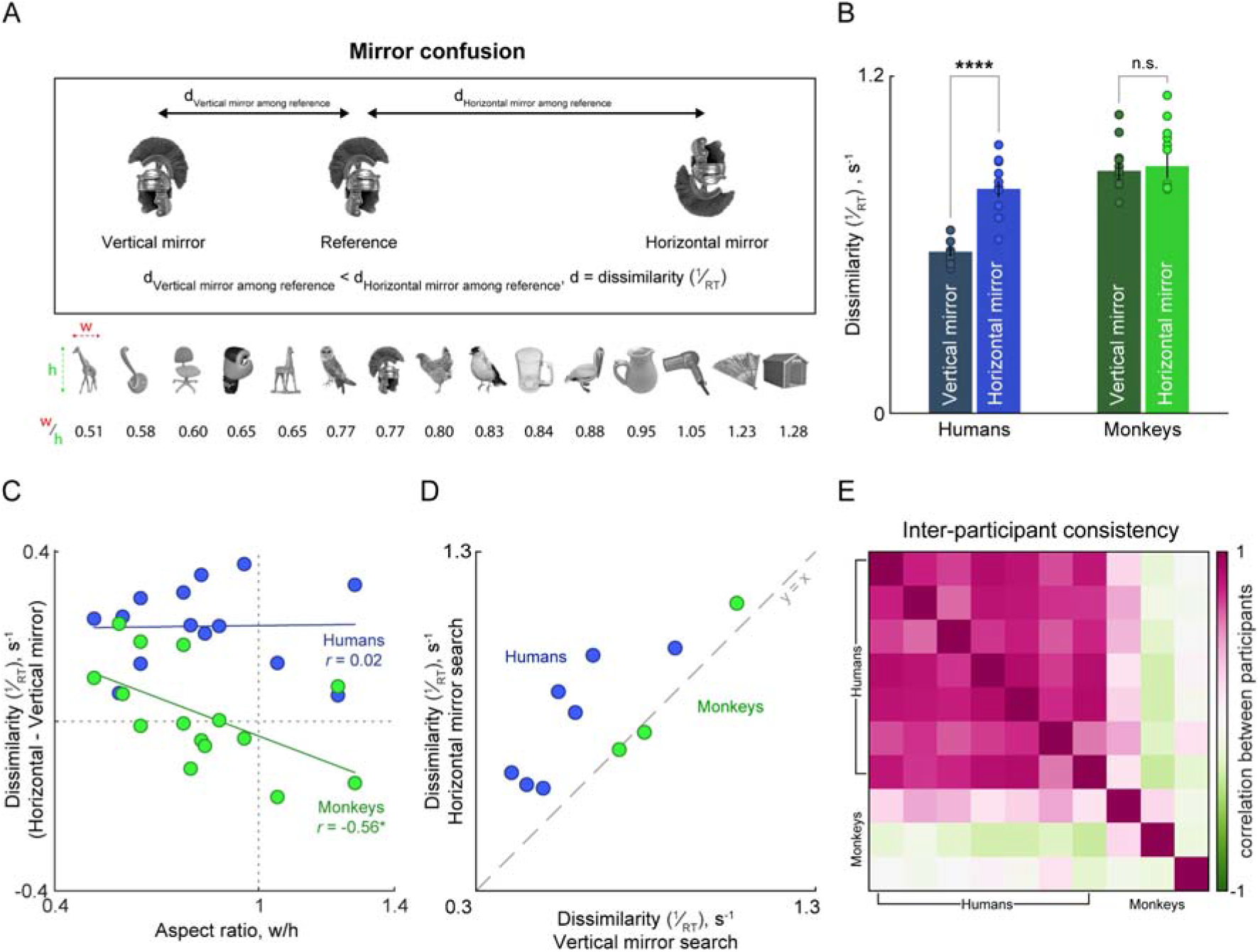
Monkeys do not show mirror confusion like humans. (A) Perception of mirror confusion of natural images in visual search for humans. Below are the 15 stimuli arranged according to the aspect ratio: calculated as the ratio of the width to the height. (B) Mean ± s.e.m. of the dissimilarities for humans and monkeys across the two types of trials, vertical mirror among reference: the target was a vertical mirror image embedded among reference displays, horizontal mirror among reference: the target was a horizontal mirror image embedded among reference displays, asterisks indicate significance of the Student’s t-test checking for differences between the means where **** is *p* < 0.0005, and n.s. is *p* > 0.05 (paired samples), data points are unique trials (averaged across all subjects and repetitions). (C) *Effect strength for each search.* Difference in dissimilarity between horizontal and vertical mirror conditions plotted against the aspect ratio of the 15 stimuli tested for humans (*blue*) and monkeys (*green*). The *r* values are the Pearson correlation and coloured lines are the best-fitting lines for each group. (D) *Effect strength for each participant.* Mean dissimilarities for vertical mirror among reference and horizontal mirror among reference pairs for each subject, 7 humans and 3 monkeys. (E) Inter-participant consistency for all pairs of participants (both human and monkey), calculated as the pairwise correlation between participants’ object dissimilarities. Large values indicate high agreement between that pair of subjects.

Humans were highly accurate during visual search involving mirror images (accuracy, mean ± s.d.: 95% ± 2%, chance is 25%) and their reaction times were also highly consistent (*r_sh_*= 0.92, *p_sh_* < 0.001, *n* = 30 search pairs). They were also more accurate on horizontal compared to vertical mirror image searches (accuracy, mean ± s.d.: 93% ± 3% for vertical mirror searches; 99% ± 1% for horizontal mirror searches; *p* < 0.0005, sign-rank test across mean accuracy across 15 objects). This effect was more pronounced when comparing the search dissimilarity, obtained by taking the reciprocal of the average response times (Figure 5B; dissimilarity, mean ± s.e.m.: 0.57 ± 0.01 s^-1^ for vertical mirror searches; 0.80 ± 0.02 s^-1^ for horizontal mirror searches; *p* < 0.00005, paired t-test across 15 natural objects). Thus, in keeping with previous observations, humans showed strong and robust vertical mirror confusion during visual search.

Next, we asked whether monkeys showed mirror confusion. Monkeys performed the task itself well above chance (accuracy, mean ± s.d.: 67% ± 6% across three monkeys; chance is 25%, see example trials in <u>Movie S1</u>), and their reaction times were also consistent (*r_sh_* = 0.81, *p_sh_* < 0.001, *n* = 30 search pairs). Their accuracy did not differ significantly between vertical and horizontal mirror searches (accuracy, mean ± s.d.: 66% ± 11% for vertical mirror searches; 69% ± 3% for horizontal mirror searches; *p* = 0.38, sign-rank test across mean accuracy across 15 objects). We also observed no significant difference in their dissimilarities for vertical vs horizontal mirror searches (Figure 5B; dissimilarity, mean ± s.e.m.: 0.86 ± 0.03 s^-1^ for vertical mirror searches; 0.88 ± 0.04 s^-1^ for horizontal mirror searches; *p* = 0.59, paired t-test across 15 natural objects). Thus, monkeys do not experience mirror confusion during visual search like humans do.

Because prior studies with monkeys have demonstrated mirror-confusion (Riopelle et al., 1964; Washburn, 1993; Rollenhagen and Olson, 2000), we dug deeper into our data to investigate if the monkey behaviour was driven by any stimulus-dependent factors. We found that monkeys do show mirror confusion for some objects but not others. The effect seemed to be strongly driven by the aspect ratio (width/height): a plot of the difference in dissimilarity (horizontal minus vertical) against the aspect ratio for each object revealed a very interesting difference in behaviour between humans and monkeys (Figure 5C). Humans showed a consistent mirror confusion independent of aspect ratio (Figure 5C, *r* = 0.02, *p* = 0.935). By stark contrast, monkeys showed a strong negative correlation (Figure 5C, *r* = -0.56, *p* = 0.029). In other words, for narrow objects they had higher dissimilarity for horizontal mirror images compared to vertical mirror images, whereas the trend was reversed for wide objects. This suggests that they were using very different features to search for oddball targets compared to humans.

To elucidate whether mirror confusion is present in individual participants, we plotted the dissimilarity for each participant for horizontal mirror searches versus vertical mirror searches (Figure 5D). Here, vertical mirror confusion would be indicated by values above the y = x line. It can be seen that all human participants showed a mirror confusion effect whereas the effect is weak or absent across the three monkeys.

Finally, to investigate how similar participants are to each other in their responses, we calculated the inter-participant consistency as before. This revealed a remarkable grouping of human and monkey participants: human participants were more similar to each other in their responses, whereas monkeys were dissimilar to humans and to each other in their responses (Figure 5E).

We conclude that monkeys do not show mirror confusion during visual search as humans do.

### Experiment 5. Do monkeys show an advantage for global shapes?

Humans are faster to report shape at the global level compared to the local level (Navon, 1977; Kimchi, 1992). In visual search, this has been demonstrated as faster responses to changes in global shape compared to local shape (Jacob and Arun, 2020), as depicted in Figure 6A. We therefore set out to investigate whether monkeys show global advantage during visual search like humans do.

**Figure 6.**
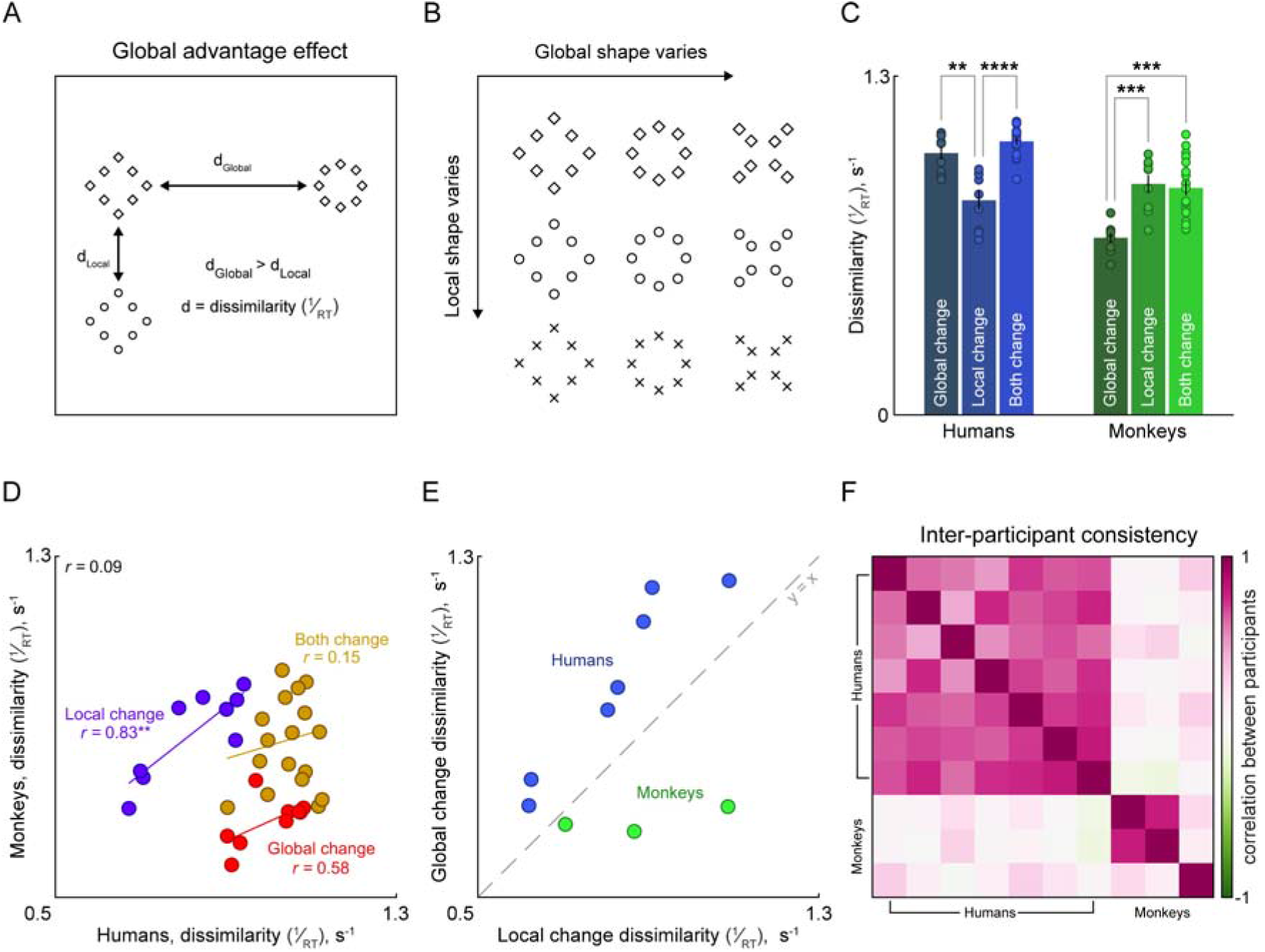
Monkeys show a local advantage unlike humans. (A) Humans show a global advantage when viewing hierarchical stimuli, such as a circle made of diamonds. Specifically, for humans, shapes differing only at the global level are perceived as more dissimilar than shapes differing only at the local level (i.e. d_Global_ > d_Local_). (B) Hierarchical stimuli used for the visual search experiment in humans and monkeys. An example video of one of the monkeys performing this task is available in <u>Movie S1</u>. (C) Dissimilarity between pairs of hierarchical stimuli during visual search, in humans (*left group, blue*) and monkeys (*right group, green*). Dissimilarity is shown separately for three groups of search pairs: pairs differing only in global shape (*left bars*), differing only in local shape (*middle bars*) and differing in both global and local shape (*right bars*). Dots centred around each bar represent average dissimilarity for each individual participant. Asterisks indicate statistical significance obtained from an unpaired t-test comparing the mean dissimilarities in each group (*n* = 9 for global change and local change, *n* = 18 for both change, ** is *p* < 0.005, *** is *p* < 0.0005, ***** is *p* < 0.00005, non-significant comparisons are not marked). (D) *Effect strength for each condition between humans and monkeys.* Average dissimilarity for each search pair in monkeys plotted against the corresponding values in humans. The overall correlation between all search pairs is shown on the top left (*r* = 0.09). Correlations for each group of pairs are also shown separately: pairs differing only in global shape (*red dots*), in local shape (*purple dots*) and differing in both global and local shape (*ochre dots*). Asterisks represent statistical significance (** is *p* < 0.005). Correlation values without an asterisk are not statistically significant. (E) Effect strength for each participant. Average dissimilarity for each search pair with a global shape difference plotted against the dissimilarity for the corresponding local shape difference for individual participants (humans, *blue dots*; monkeys, *green dots*). It can be seen that all humans find global shape differences consistently larger than local shape differences, whereas all monkeys show exactly the opposite pattern. (F) Inter-participant consistency for all pairs of participants (both human and monkey), calculated as the pairwise correlation between participants’ object dissimilarities. Large values indicate high agreement between that pair of subjects.

To this end, we created hierarchical shapes with either a diamond, circle, or ‘x’ shape at the global, local or both levels (Figure 6B). We created search arrays involving all pairs of these shapes: these search arrays could therefore be divided into those that contained only a change in global shape, a change in local shape, or both global and local shape changes.

Humans were highly accurate on this task (accuracy, mean ± s.d.: 96% ± 1%) and their reaction times were also highly consistent (*r_sh_*= 0.93, *p_sh_* < 0.001, *n* = 36 search pairs). They were also more accurate to find global compared to local shape changes (accuracy, mean ± s.d.: 97% ± 02% for global change searches; 91% ± 04% for local change searches; *p* < 0.05 using sign-rank test comparing 9 global vs local change searches). Humans showed significantly smaller dissimilarities for searches with local compared to global shape differences, confirming that they are more sensitive to global shape changes (Figure 6C; dissimilarity, mean ± s.d.: 1.00 ± 0.02 s^-1^ for searches with only global change; 0.82 ± 0.03 s^-1^ for searches with only local change, *p* < 0.0005, paired t-test for global vs local change across 9 searches). Interestingly, searches with both global and local change were even faster than searches involving only one type of change, suggesting that humans integrate both local and global shape during search (Figure 6C; dissimilarity, mean ± s.e.m.: 1.05 ± 0.06 s^-1^ for searches with both change; *p* < 0.00005 comparing both change with local-only changes and *p* = 0.09 comparing both change with global-only changes, unpaired t-test). Thus, consistent with previous studies, humans showed a robust global advantage during visual search.

Next, we asked whether monkeys showed a global advantage. Monkeys performed poorly on visual search involving these hierarchical stimuli (accuracy, mean ± s.d.: 56% ± 15% across three monkeys; chance is 25%) but their reaction times were highly consistent (*r_sh_* = 0.85, *p_sh_* < 0.001, *n* = 30 search pairs). They also showed an accuracy difference between global and local shape changes in favour of local shape changes (accuracy, mean ± s.d.: 31% ± 01% for global change searches; 66% ± 19% for local change searches; *p* = 0.003 using sign-rank test comparing 9 global vs local change searches).

However, monkeys showed a dramatically different pattern from humans. This is illustrated by one of the monkeys doing the task in <u>Movie S1</u>. Monkeys showed a significantly smaller dissimilarity for global shape changes compared to local shape changes (Figure 6C; dissimilarity, mean ± s.e.m.: 0.67 ± 0.01 s^-1^ for searches with only global change; 0.88 ± 0.03 s^-1^ for searches with only local change; *p* < 0.0005, paired t-tests comparing 9 global and local searches). Even for searches with both global and local changes, monkeys’ performance was better than searches with global-only changes but no better than their performance on searches with local-only changes (Figure 6C; dissimilarity, mean ± s.e.m.: 0.87 ± 0.02 s^-1^ for searches with both global and local change; *p* < 0.0005 for both vs global-only changes; *p* = 0.73 for both vs local-only changes, unpaired t-test). Thus, monkeys showed markedly smaller sensitivity to changes in global shape and were more sensitive to local shape changes, exactly opposite to humans.

To investigate these differences between monkeys and humans further, we plotted the search dissimilarity in monkeys for each search condition against the corresponding values obtained from human participants (Figure 6D). This revealed very interesting patterns. There was no overall correlation between human and monkey dissimilarities across all 36 search pairs tested (Figure 6D, *r* = 0.09, *p* = 0.59). Interestingly, human and monkey dissimilarities were correlated only for searches involving local-only shape differences (Figure 6D; *r* = 0.83, *p* < 0.005 for local-shape changes) but not for global-only searches or both global & local change searches (Figure 6D; *r* = 0.58, *p* = 0.09 for global-only searches; *r* = 0.15, *p* = 0.54 for global & local searches). The poor correlation for global-change-only searches might be due to poor accuracy in those searches for monkeys. Furthermore, dissimilarities for both-change searches also do not agree as both humans and monkeys appear to use different information to make their decisions. Thus, monkeys seem to prioritize information at the global and local levels differently from humans when both types of information is present, but process information at the local level similarly to humans.

To elucidate individual differences, we plotted the average dissimilarity for each participant for global changes against the corresponding dissimilarity for local changes (Figure 6E). This too revealed a remarkable difference: all human participants showed greater dissimilarity for global changes, whereas all monkeys showed greater dissimilarity for local changes.

This striking difference between humans and monkeys was also evident upon calculating inter-participant consistency as before. Here, we found that human participants were highly consistent with each other and behaved quite differently from monkeys. Likewise, all three monkeys were more consistent with each other than with human participants.

We conclude that monkeys show a local advantage, unlike humans.

### Experiment 6: Why do monkeys show a local advantage?

The local advantage observed in monkeys could be due to two reasons. One possibility is that monkeys cannot integrate spatially disconnected local elements into a larger global shape. Alternatively, they might prioritize information at smaller spatial scales compared to humans. To distinguish between these possibilities, we performed an additional experiment. We took the same shapes as used earlier but added connecting gray lines between the local shapes so that the global shape was evident without the requirement of grouping (Figure 7A).

**Figure 7:**
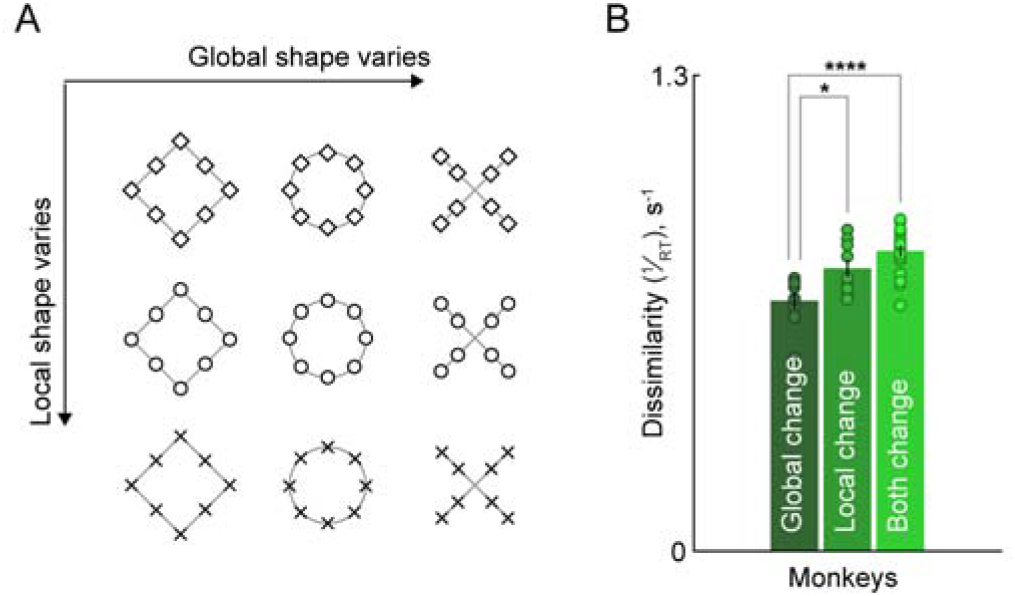
Monkeys show a local advantage even for connected shapes. (A) Hierarchical stimuli used for this experiment. Here, we connected local elements within each global shape with a gray line to facilitate grouping into a global shape. (B) Avergage dissimilarity between pairs of hierarchical stimuli during visual search in monkeys, shown separately for three groups of search pairs: pairs differing only in global shape (*left bars*), differing only in local shape (*middle bars*) and differing in both global and local shape (*right bars*). Dots centred around each bar represent average dissimilarity for each individual participant. Asterisks indicate statistical significance obtained from an unpaired t-test comparing the mean dissimilarities in each group (*n* = 9 for global change and local change, *n* = 18 for both change, * is *p* < 0.05, **** is *p* < 0.00005, non-significant comparisons are not marked).

All three monkeys were well above chance on this task (accuracy, mean ± s.d.: 59% ± 12% across three monkeys, chance is 25%) and their reaction times were consistent (*r_sh_* = 0.80, *p_sh_* < 0.001, *n* = 30 search pairs). As in Experiment 5, they also showed an accuracy difference between global and local shape changes in favour of local shape changes (accuracy, mean ± s.d.: 43% ± 07% for global change searches; 62% ± 13% for local change searches; *p* = 0.02 using sign-rank test comparing 9 global vs local change searches).

Here too, monkeys showed a larger dissimilarity for local changes compared to global changes (Figure 7B, mean ± s.e.m.: 0.68 ± 0.02 s^-1^ for searches with only global change; 0.77 ± 0.02 s^-1^ for searches with only local change; *p* < 0.05, paired t-test across 9 search pairs). Even for pairs differing in both local and global shape, monkeys behaved as if they only processed local shape: the dissimilarity for shapes differing at both levels was larger than global-only changes, but similar to local-only changes (average dissimilarity, mean ± s.e.m.: 0.81 ± 0.01 s^-1^ across 18 searches with both global and local change; *p* = 0.091 for local change vs both change; *p* < 0.00005 for global change vs both change; unpaired t-test).

Thus, local advantage in monkeys is present even when local elements are explicitly connected together, suggesting that their local advantage comes from a higher priority for local shape rather than an inability to group local elements. It would be interesting to explore these differences further in future work.

## GENERAL DISCUSSION

Here, we compared perception in monkeys and humans by testing them on a wide range of high-level perceptual phenomena. Monkeys showed similar coarse object representations, Weber’s law and rapid amodal completion just like humans. However, they did not show mirror confusion or a global advantage, unlike humans. These findings are summarized below in Table 1. Our results reveal both qualitative similarities and differences between human and monkey perception and ultimately provide for an unbiased and nuanced evaluation of macaque monkeys as a model for human vision. Below we discuss these findings in relation to the existing literature.

**Table 1:** Summary of perceptual phenomena observed in monkeys vs humans.

| Perceptual Phenomena | Monkeys | Humans |
| --- | --- | --- |
| Object Relations | Yes | Yes |
| Weber's Law | Yes | Yes |
| Amodal Completion | Yes | Yes |
| Mirror Confusion | No | Yes |
| Global Advantage | No | Yes |

### Relation to previous approaches

There is a long and rich literature of studies on visual perception in non-human primates and comparisons to human vision (Matsuno and Fujita, 2009). Comparative studies have included diverse non-human primate species including marmosets, baboons, macaques and even chimpanzees. Because of the difficulty of obtaining any kind of report from animals, each study usually has only tested one specific aspect of visual perception in non-human primates. These tests often involve extensive training of animals on complex tasks. Such extensive training could potentially modify the underlying visual representations or the decision-making process.

Our study represents a novel alternative to study animal perception. We trained animals on an oddball search task that could be performed on any set of images. We then selected a range of perceptual phenomena that could be tested by choosing different sets of images. This approach was successfully used recently to investigate deep networks (Jacob et al., 2021b) and proposed as a general framework where testing different ‘signatures’ allows the comparison of cognitive processes across multiple species (Taylor et al., 2022). For animal cognition, our approach has the major advantage that animals need not be retrained each time to test each additional aspect of perception. As a result, our approach also avoids the danger of extensive training modifying the underlying representations.

An interesting alternative approach to understanding perception in animals is a free-choice paradigm, pioneered by Logothetis and colleagues to study the neural correlates of subjective percepts in monkeys (Logothetis and Schall, 1989; Leopold and Logothetis, 1996). In these studies, animals were first trained to report unambiguous stimuli first where only the correct response is rewarded, and then presented in a small fraction of trials containing ambiguous binocular rivalry stimuli where both responses are rewarded. The animals’ response in the ambiguous trials were highly systematic and were taken as indicative of their subjective percept. Such free choice paradigms could be extended to other aspects of perception such as those tested in our study. For instance, to test mirror confusion, monkeys could be trained on a same-different task involving pairs of completely different images where they are rewarded only for making correct responses and then presented in a small fraction of trials with vertical-mirror and horizontal-mirror pairs of images where either response would be rewarded. If monkeys made more “same” responses for vertical-mirror compared to horizontal-mirror pairs, this would confirm that they experience mirror confusion. We considered such free-choice paradigms but decided instead to use visual search for several reasons. First, free-choice paradigms by design will only have a small fraction of trials of interest, limiting our ability to test many aspects of perception. Second, free choice paradigms work best if all trials are similar in difficulty, but even small variations in difficulty could make animals change their response strategies when they encounter hard trials. Third, free-choice paradigms also involve extensive training to achieve extremely high levels of performance. In our experience, monkeys are highly sensitive to task difficulty and can often switch their task strategy fluidly from following a rule to making random choice responses. By contrast, visual search in our experience was a much more natural task for monkeys to learn and can be performed at a high throughput. Importantly, we were easily able to elicit large variations in response times without affecting the overall accuracy and reward rate, thereby reducing the animals’ incentive to switch task strategies and enabling them to operate according to a stable decision rule throughout the session. We therefore propose that visual search could be a highly effective task to study animal perception across multiple species.

### Relation of each finding to previous studies

Below we discuss each individual finding in our study with previous studies. Our first finding is that monkeys and humans have similar object representations in visual search. In other words, searches that were hard for humans were hard for monkeys as well (Figure 2). This finding is consistent with a number of previous studies showing similar object categorization behaviour (Rajalingham et al., 2015, 2018), and similar object representations at the neural level (Kriegeskorte et al., 2008b) in monkeys and humans. In fact, neural representations in monkey IT cortex are highly predictive of visual search dissimilarity in humans (Sripati and Olson, 2010; Zhivago and Arun, 2014). These studies together suggest that macaque monkeys and humans have similar coarse object representations, or at the very least, use similar features when responding to natural objects.

Our second finding is that monkeys show Weber’s law for luminance, just like humans (Figure 3). Weber’s law has been demonstrated in humans for luminance discrimination for intensity values within a broad range (Hecht, 1924; Pramod and Arun, 2014), line length (Ono, 1967; Pramod and Arun, 2014) and sound pressure levels (Pardo-Vazquez et al., 2019). While Weber’s Law has been demonstrated in monkeys for numerosity (Nieder and Miller, 2003; Jordan and Brannon, 2006) and line length (Tudusciuc and Nieder, 2010), to our knowledge our study is the first to demonstrate Weber’s law for luminance discrimination in monkeys.

Our third finding is that monkeys experience rapid and automatic amodal completion in visual search, just like humans (Figure 4). There is only one similar study where monkeys trained to categorize line length respond to an occluded line as being longer (Fujita, 2001). By contrast, we demonstrate using a wide range of abstract shapes that monkeys find it difficult to find an occluded target among likely completions compared to a mosaic target among likely completions, just like humans do (Rensink and Enns, 1998; Cherian and Arun, 2024). Our study is the first to demonstrate that monkeys experience rapid, automatic amodal completion in the same way as humans.

Our fourth finding is that monkeys do not show consistent mirror confusion, unlike humans (Figure 5). This disagrees with previous reports of mirror confusion in monkeys both in behaviour (Riopelle et al., 1964; Washburn, 1993) as well as in their inferior temporal cortex (Rollenhagen and Olson, 2000). There could be several reasons for this discrepancy. The task in the two previous studies involved training monkeys to discriminate between two mirror images that were either horizontal mirror reflections or vertical mirror reflection, and the images used were letters or symbols. By contrast, monkeys in our study did not receive any special shape discrimination training and were tested on images of real animate and inanimate objects. Reconciling these differences will involve testing monkeys on both natural objects as well as abstract shapes. Our finding also stands in contrast with the observation of mirror confusion in single neurons in object-selective and face-selective cortex in macaque monkeys (Rollenhagen and Olson, 2000; Freiwald and Tsao, 2010). We speculate that monkeys might have such representations but might not necessarily use them to guide visual search. These are interesting possibilities for future study.

Our final finding is that monkeys show a local advantage while processing stimuli containing global and local shape, unlike humans who show a global advantage (Figure 6). This local advantage persisted even when we connected the local elements, suggesting that the local advantage may not arise from differences in perceptual grouping. It is possible that monkeys would show a global advantage for other variations of these shapes, such as solid silhouettes or geometric shapes (Sablé-Meyer et al., 2021). There is mixed evidence in the literature regarding this effect in non-human primates, with many studies reporting a local advantage and some a global advantage. Baboons and capuchin monkeys trained on shape matching tasks show local advantage (Deruelle and Fagot, 1997; Fagot and Deruelle, 1997; Spinozzi et al., 2006). Interestingly, chimpanzees trained on oddball search have a local advantage during visual search but a global advantage when the local elements are connected (Fagot and Tomonaga, 1999). By contrast, macaque monkeys continued to show a local advantage in our study even when the local elements are connected. When tested on the same stimuli and task, chimpanzees showed a global advantage, but macaque monkeys showed a local advantage (Hopkins and Washburn, 2002). Thus, there could be differences even between non-human primates. Our findings are discordant with previous studies showing that macaque monkeys trained on shape matching have a global advantage (Tanaka and Fujita, 2000) and with a report of global advantage present in the macaque high-level visual cortex (Sripati and Olson, 2009). Once again, we surmise that perhaps macaques have such representations but do not necessarily use them for visual search. Evaluating this possibility will require testing for global/local advantage at the behavioural and neural levels simultaneously.

### Limitations of the present study

Although we have done our best to perform a comprehensive comparison of several perceptual phenomena between monkeys and humans, our study has several limitations.

First, we were only able to test these phenomena on three monkeys, because training monkeys on these tasks is extremely challenging, and requires months of focused training, because of which it was simply not feasible for us to test larger numbers of animals. However, we consider it unlikely that our results would not generalize, because many of the tested perceptual phenomena were robust across our monkeys, and are highly robust across humans. There was also striking similarity between the three monkeys, which would not have been observed had there been large inter-individual differences. Even if our results did not generalize, the fact remains that a subpopulation of monkeys do show similarities and differences as observed here. Testing these phenomena on larger populations of animals will require installing automated training stations in group enclosures where many animals can be tested simultaneously.

Second, the perceptual phenomena tested in this study are based on small numbers of images and conditions that again might not generalize when repeated on a larger set of images/conditions. While this is possible in principle, it was simply not possible for us to test a wide range of phenomena with a wide range of images, in the interest of having compact experiments that could be completed within one testing session. Testing a larger set of conditions would require multiple sessions for each animal, which would require counterbalancing sessions and introduce potential practice effects. However, we consider it unlikely that our results would not generalize across a wider set of conditions/images, because these effects are highly robust when tested in humans with relatively little inter-individual variability.

Third, some of the perceptual phenomena tested in this study are based on testing natural objects, that are likely to be familiar to humans but not necessarily to our monkeys. As a result, the results observed could potentially be explained by familiarity to these images in humans but not in monkeys. We consider this unlikely, for several reasons. Our monkeys were housed in natural arenas where they encountered many natural objects, although not necessarily the images used in our experiments. While our human participants might have indeed been familiar with some objects, their responses during visual search may not have been driven by familiarity, particularly since their task involved searching for oddball targets without specifying their features. Indeed, oddball search shows only subtle effects of familiarity, as evidenced by highly correlated performance between readers and nonreaders on letter searches (Agrawal et al., 2019, 2022). Understanding how familiarity shapes object representations in humans and monkeys would require careful testing with increasingly familiar objects and potentially with arbitrary shapes that are unfamiliar to both species. These are important considerations for future studies of cross-species comparisons.

### Implications for macaques as a model system for vision

Our study represents a first comprehensive evaluation of visual perception in monkeys on multiple perceptual phenomena and revealed important similarities as well as differences between monkey and human vision. The qualitative similarities with human vision reconfirms that macaque monkeys are indeed good models. However, the qualitative differences observed in our study also demonstrate important limitations of macaque monkeys as a model for human vision. However, we must be cautious in interpreting these similarities and differences, because of the important distinction between performance and competence (Firestone, 2020). In other words, macaque monkeys and humans may have similar visual representations, but their overt behaviours might be different due to the way in which they utilize these representations to perform tasks. This could explain the contradictory findings between our study and previous work: for instance, there is mirror confusion and global advantage in the high-level visual areas of macaque monkeys (Rollenhagen and Olson, 2000; Sripati and Olson, 2010), but we did not observe these effects in our study in behaviour. Of course, these results come from testing neural representations in one group of macaques and testing behaviour in another, so reconciling these differences will require comparing these phenomena in the same animals.

### Conclusions

Our study represents a novel and general approach to characterize a variety of visual systems, similar to the signature testing approach proposed for cognition (Taylor et al., 2022). We propose that visual search can be a powerful paradigm to test a variety of animals since is a natural, ecologically valid task with an objective performance measure. Visual search dissimilarities can be directly compared to neural dissimilarities, enabling comparing neural representations (Sripati and Olson, 2010; Zhivago and Arun, 2014). These can also be compared with dissimilarities in neural network activations (Jacob et al., 2021b) or computer vision algorithms (Pramod and Arun, 2016b). This enables powerful comparisons between a variety of visual systems for core perceptual phenomena, which we believe will yield fundamental insights into the nature of vision and perception.

## MATERIALS AND METHODS

All human experiments were performed according to a protocol approved by the Institutional Human Ethics Committee at the Indian Institute of Science (IHEC # 10/20.07.2022). All animal experiments were performed according to a protocol approved by the Institutional Animal Ethics Committee of the Indian Institute of Science and by the Committee for the Purpose of Control and Supervision of Experiments on Animals, Government of India (V-11011(3)/15/2020-CPCSEA-DADF).

### Humans

All participants were recruited from the Indian Institute of Science community, had normal or corrected-to-normal vision and were monetarily compensated. In addition, they were unaware of the experimental goals prior to completing the tasks. Experiments 2, 4 and 5 were conducted on the same setup as used to test monkeys, whereas Experiments 1 & 3 were obtained from previous studies (see below).

We recruited 16 participants for Experiment 2 (6 female, 10 male; ages mean ± s.d.: 26.1 ± 3.6 years), 7 participants for Experiment 4 (4 female, 3 male; ages mean ± s.d.: 26.7 ± 2.2 years) and 7 participants for Experiment 5 (4 female, 3 male; ages mean ± s.d.: 23.7 ± 3.0 years). For Experiment 1 (12 participants; 3 female; aged 20-30 years) & Experiment 3 (7 female, 9 male; ages mean ± s.d.: 27.5 ± 3.6 years), we used previously collected data from human participants performing a similar oddball search task (Mohan and Arun, 2012; Cherian and Arun, 2024).

### Monkeys

Three male adult bonnet macaques (*macaca radiata*; all aged ∼8 years, laboratory designations: *Di*, M1; *Ju*, M2; and *Co*, M3) were tested in this study. The monkeys were under fluid restriction during data collection but had unlimited access to food. These monkeys were initially trained on same-different tasks and fixation tasks previously (Jacob et al., 2021a) and were subsequently trained to perform the oddball search task for this study, as follows.

Monkeys were initially trained by showing them oddball search arrays containing colored circles, with the same colour pair as the target and distractor. Since monkeys could simply learn to associate a colour (not the oddball) with a reward, subsequent stages of training involved including additional colour pairs. Once the monkey’s performance across >10 color pairs reached a high level of accuracy, we intermixed all colours as target or distractor. Monkey performance did not change when we switched to natural objects as targets and distractors, and did not need any additional familiarization or training with the experimental stimuli. The entire process of training took about 2-3 weeks.

### Apparatus

We used a touchscreen-based behaviour station for our data collection for monkeys (Jacob et al., 2021a). Briefly, monkeys sat at a juice spout in front of a touchscreen (15” capacitive touchscreen, 1366 x 768 pixels, Elo Touch Solutions Inc. Model 1593L RevB) at an eye-to-screen distance of 23 cm. For humans, we replicated the setup on a table such that humans could sit comfortably in front of the screen with the same eye-to-screen distance as monkeys. All experiments were presented using NIMH MonkeyLogic software (Hwang et al., 2019) running on MATLAB (R2020b, The Mathworks Inc.). Monkeys were rewarded with juice delivered through a custom reward circuit and peristaltic pump that allowed for precise dosing.

### Task

Both humans and monkeys performed the same oddball visual search task (Figure 1A). At the start of the trial, a yellow button (a circle with a radius of 4°, degree of visual angle) appeared on the screen against a black background. Subjects had to hold this button to initiate the trial within 10 seconds. After this, four stimuli (three identical items and one oddball item) appeared on the screen (see Figure 1B and 1C), arranged circularly around the left side of the hold button. Stimuli were presented 15° away from the hold button centre and 36° apart from each other. For Experiment 2, stimuli appeared in a row aligned to the screen centre and spaced 9° apart. Participants had to touch any target to complete the trial; touches to the oddball target were considered a correct response. On making a correct response, a rising auditory tone was played, and monkeys were rewarded with juice. Correct trials were followed by a short inter-trial interval (200 ms) and errors by a slightly longer inter-trial interval (500 ms).

For two experiments (Experiments 1 and 3), human data was obtained from previous studies from our lab where participants also performed a similar oddball visual search task (Figure 1D). Briefly, participants were seated in front a monitor and were presented with either a 16-item search array (4-by-4 grid, Experiment 1) or a 12-item search array (3-by-4 grid, Experiment 3), containing an oddball item among identical distractors. Participants had to indicate using a keypress the side on which the oddball item appeared (left vs right).

### Stimuli and search pairs

All stimuli were grayscale and scaled to subtend 5° along their longest dimension. Across the duration of each experiment, the target location was sampled equally across each of the four possible choices in the search array. Each set of unique trials (i.e., search pairs) were formed into a block when running the task. Within each block the trials were presented in random order. Thus, repetitions of unique trials were collected as blocks. Due to an unforeseen issue in the applied settings in NIMH MonkeyLogic, for some experiments (Experiments 1, 3 and 4 in monkeys), all conditions did not have equal repetitions, and we performed all analyses using all available trials for each condition. Environmental illumination was present via roof mounted lamps. Any deviations from the preceding statements are mentioned explicitly for each experiment.

#### Experiment 1 – Perceptual organization

We selected four animate and four inanimate object images (Figure 2A). Unique search pairs consisted of all ^8^C_2_ = 28 image pairs. For each unique search pair, we created two search arrays with either item as the target and the other as the distractor for a total of 56 search pairs. Each search pair was presented 12-20 times to each monkey. This was compared to human data collected for the same stimuli and image pairs in a previously published study (Mohan and Arun, 2012), where each search pair was presented twice to each participant.

#### Experiment 2 – Weber’s Law for luminance

To test perceptual discriminability of luminance, we created circles of size 2° each and set the luminance levels based on a 16-bit scale (a.u.). Actual luminance values (in cd/m^2^) were measured post-hoc using a SpectraScan PR655 spectroradiometer (PhotoResearch Inc.). In each trial, the distractors’ luminance was either low-intensity (16384 a.u., 8.8 cd/m^2^) or high-intensity (25600 a.u., 24.33 cd/m^2^) and the targets’ luminance was greater than the distractor by a fixed difference in the range of 500 to 8000 a.u. (10-linearly spaced steps, ranging from 9.13 to 21.66 cd/m^2^ for low-intensity distractors and from 24.93 to 45.30 cd/m^2^ for high-intensity distractors). For each distractor luminance level, there were 10 trials with each of the targets for a total of 20 unique search pairs. Each search pair was presented 16 times to monkeys and 12 times to humans. The experiment was blocked to ensure that trials for a given distractor luminance level appeared together. We turned off the room lights to promote task performance.

#### Experiment 3 – Amodal completion

As in Experiment 1, we obtained the stimuli and the relevant human data from a previous study (Cherian and Arun, 2024). To test amodal completion in monkeys, we selected 10 shapes (Figure 3A) and created 3 display conditions for each. In the ‘occluded’ displays an occluder blocked the top left quadrant of the shape from view whereas in the ‘likely’ and ‘mosaic’ displays the occluder was positioned so as to not occlude any portion of the shape. While the ‘likely’ displays showed the whole shape intact, in the ’mosaic’ displays the top left quadrant of the shape was cropped out. For each shape we prepared two search pairs, with the ‘likely’ display always as the distractor and the ‘occluded’ or ‘mosaic’ display as the target. Thus, we created a total of 20 unique search pairs. Each search pair was presented 15-24 times to each monkey with varying target location and twice for each human participant.

#### Experiment 4 – Mirror confusion

We selected images of 15 natural objects from the BOSS dataset (Brodeur et al., 2014). We labelled these images as ‘reference’ and created ‘vertical’ and ‘horizontal’ mirror images of each. Each search pair consisted of the ‘reference’ image as the distractor and either the ‘vertical’ or ‘horizontal’ mirror image as the target, for a total of 30 unique search pairs. All search pairs were presented 18-29 times to each monkey and 8 times to each human participant.

#### Experiment 5 – Global advantage

Stimuli were obtained from a previous study (Jacob and Arun, 2020). Three basic shapes: ‘diamond’, ‘circle’ and ‘x’ were applied on both a global and local scale to create a set of 9 hierarchical shapes with all possible mixtures (Figure 6B). For any pair of these shapes either the global, local or both shapes were different. We used all 36 unique pairs i.e., the 9 global same and local different (GSLD), 9 global different and local same (GDLS) and 18 global different and local different (GDLD) pairs. For each search pair we created two search arrays with either item as the target and the other as the distractor for a total of 72 total search pairs. Monkeys were presented with each of the search pairs 8 times whereas humans were presented with 4 repetitions.

#### Experiment 6 – Global advantage with connected lines

Stimuli were the same as Experiment 5 except that we connected the local elements with a gray line with the same thickness as the lines making the local shapes (Figure 7A). We chose a grey colour for the connecting line to ensure local shapes remained individually distinguishable. All other experimental details were identical to Experiment 5.

### Data analyses

We analysed all data through custom scripts and functions in MATLAB. Only trials where humans or monkeys gave a valid response (correct or wrong) were included for all further analyses.

#### Response consistency

To evaluate response time consistency for humans and monkeys, we first collated the response times across all participants (either human or monkey). For each search pair, outliers more than 3 scaled median absolute deviations from the median were removed for each search pair. Doing so improved the response consistency overall for both humans and monkeys. We confirmed that all results were qualitatively similar even without this step. We then calculated the response consistency as the split-half correlation as the average Pearson’s correlation coefficient between 1000 random splits (*r_sh_*, with repeated sampling). The statistical significance of this correlation (*p_sh_*) was determined as follows:

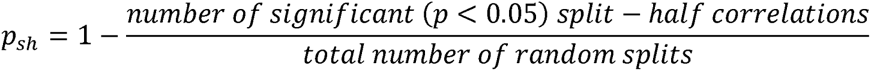

#### Measuring perceptual dissimilarities

We analysed only response times (RT) from correct trials, averaged them across repetitions and took the reciprocal (1/RT) as a measure of perceptual dissimilarity. 1/RT behaves like a distance metric, varies linearly with feature differences, and can be interpreted as the underlying evidence accumulation signal driving search decisions (Arun, 2012; Pramod and Arun, 2014; Sunder and Arun, 2016).

#### Inter-participant consistency

To compare inter-individual differences across humans and monkeys, we calculated the correlation distance between every pair of participants as follows:

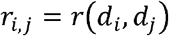

Where *r_i_,_j_* is the Pearson’s correlation between the set of dissimilarities, *d* (averaged across repetitions), from subjects *i* and *j*. A large correlation value implies that the two participants had a good match between their responses and vice-versa.

#### Sigmoid curve fitting

In order to compare psychometric curves, we fit a sigmoid by scaling and shifting the normal cumulative distribution function (CDF, ‘*normcdf*’ in MATLAB) as follows:

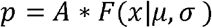

Where *p* is the predicted accuracy, *A* is the scaling factor, and *F* is the normal CDF with parameters: μ and σ *x* is the luminance difference and the values of *A* μ, and μ were estimated using non-linear least squares fitting (‘*nlinfit*’ in MATLAB), with initial guess as max accuracy, and mean and standard deviation of the luminance levels, respectively.

## AUTHOR CONTRIBUTIONS

TC, GJ and SPA designed the experiments; TC and GJ collected the data; TC and SPA analysed and interpreted the data and wrote the manuscript with inputs from GJ.

## ACKNOWLEDGEMENTS

We thank Aditya Murthy, Supratim Ray, and Sridharan Devarajan for their valuable inputs during yearly reviews. We thank Ankan Biswas and Divya Gulati for their help with the luminance measurements. We are grateful to Ramesh Venkatarayappa, Thirumala M, KV Ashok Kumar, and Ravi Kumar at the Primate Research Laboratory of the Indian Institute of Science for their excellent support with animal care.

## DATA AVAILABILITY

All data and code required to reproduce the results are available on OSF at https://osf.io/63m5f.

